# Polymeric perfluorocarbon nanoemulsions are ultrasound-activated wireless drug infusion catheters

**DOI:** 10.1101/315044

**Authors:** Q Zhong, BC Yoon, M Aryal, JB Wang, A Karthik, RD Airan

**Author notes:** These authors contributed equally to this work (Q Zhong, BC Yoon). To whom correspondence should be addressed: Raag Airan MD PhD, Departments of Radiology (Neuroradiology) and, by courtesy, Materials Science & Engineering Stanford University.

## Abstract

Catheter-based intra-arterial drug therapies have proven effective for a range of oncologic, neurologic, and cardiovascular applications. However, these procedures are limited by their invasiveness, as well as the relatively broad drug spatial distribution that is achievable with selective arterial catheterization. The ideal technique for local pharmacotherapy would be noninvasive and would flexibly deliver a given drug to any region of the body. Combining polymeric perfluorocarbon nanoemulsions with existent clinical focused ultrasound systems could in principle enable noninvasive targeted drug delivery, but it has not been clear whether these nanoparticles could provide the necessary drug loading, stability, and generalizability across a range of drugs to meet these needs, beyond a few niche applications. Here, we directly address all of those challenges and fully develop polymeric perfluorocarbon nanoemulsions into a generalized platform for ultrasound-targeted drug delivery with high potential for clinical translation. We demonstrate that a wide variety of drugs may be effectively uncaged with ultrasound using these nanoparticles, with drug loading increasing with hydrophobicity. We also set the stage for clinical translation by delineating production protocols that hew to clinical standards and yield stable and optimized ultrasound-activated drug-loaded nanoemulsions. Finally, as a new potential clinical application for these nanoemulsions, we exhibit their *in vivo* efficacy and performance for cardiovascular applications, by achieving local vasodilation in the highest flow vessel of the body, the aorta. This work establishes the power of polymeric perfluorocarbon nanoemulsions as a clinically-translatable platform for effective noninvasive ultrasonic drug uncaging for myriad targets in the brain and body.

---

Many clinical therapies utilize intra-arterial catheter infusion of a drug^1–3^ to maximize its therapeutic effect at the target, while minimizing side effects due to drug action in the non-targeted organ and body. However, these catheter-based therapies cannot necessarily achieve the desired spatial precision for a given case, due to limitations in the ability to reliably direct a catheter into a target small artery. Additionally, catheter-based therapies are invasive interventions that carry additional risks of vascular injury as well as of radiation as they are usually guided by real-time fluoroscopy. Ideally, local pharmacotherapy would be achieved noninvasively and with image-guidance that does not involve ionizing radiation, such as with optical,^4^ ultrasound,^5^ or MRI^6^ based methods. An ultrasound-gated mechanism of action would be particularly useful, given the availability of clinical MRI- or optically-guided focused ultrasound systems that may sonicate nearly any region of the body with millimetric spatial resolution, especially for lower intensity ultrasound applications.^7^

Importantly, several ultrasound-sensitive drug delivery systems have been described. These technologies may use nano- or micro-scale drug carriers that release a drug after ultrasound raises the *in situ* temperature,^8,9^ activates a ‘sonosensitizer’,^10^ or applies a sufficient peak intensity or pressure^5,11^. While high-intensity continuous wave ultrasound (for temperature-gated systems) may be difficult to achieve stably in certain regions of the body,^12^ raising the peak pressure or intensity necessitates only short bursts of ultrasound that are more straightforward to achieve *in situ*.

Perfluorocarbon nanoemulsions offer several features that are optimal for this application of noninvasive localized drug delivery: an intensity/pressure-gated drug release that should be generally applicable across a range of drugs. Indeed, such nanoemulsions have been used in preliminary proof-of-concept studies for delivery of chemotherapeutics to tumors^13,14^ or for delivering the anesthetic propofol to the brain^15^ following drug release induced by short pulses of sonication of a sufficient intensity. However, important questions have persisted as to whether these nanoemulsions could achieve the drug loading, generalizability of drug encapsulation, and stability^13^ necessary for applications beyond those few niche, proof-of-concept studies. Additionally, it has been unclear whether these particles could be produced in a manner that would enable eventual clinical translation. Finally, given the relatively large size of these nanoparticles that limits their penetration into organs, the ultrasound-induced drug release would occur in the blood pool, and doubts have existed as to whether intravascular drug release from these particles would truly result in a localized drug action. Here, we fully develop this system to address each of these open questions and explicitly demonstrate the versatility of this platform for ultrasonic uncaging of a variety of drugs, with production methods, *in vivo* biodistribution, and pharmacokinetics that enable clinical translation. Further, we demonstrate that this technology can achieve drug release that is localized in space and time to the ultrasound field, and enable local cardiovascular modulation in the highest flow vessel of the body, the aorta. Together, these results indicate that noninvasive ultrasound-induced intravascular nanoparticle uncaging is indeed a viable and readily translatable system for local pharmacotherapy for a range of organ systems.

## Results

### Moving Towards Clinical Applicability and Translation

To enable eventual *in vivo* and clinical applications, we first developed methods for producing polymeric perfluorocarbon nanoemulsions that are scalable and meet clinical standards of sterility (Fig. 1a). We focused on perfluoropentane (PFP) as the choice of perfluorocarbon to optimize between the relatively increased volatility of shorter chain compounds like perfluorobutane and the potential decreased ultrasound responsiveness of longer chain compounds like perfluorohexane. Additionally, given that PFP is currently in FDA-approved clinical trials for other applications,^16^ its inclusion would help to lower barriers to eventual translation. We targeted a median Z-average diameter of 400-450 nm, calculated so that during uncaging the nanoparticles would be at most half of a capillary diameter, given prior theory and evidence that similar particles increase their diameter up to a maximum of 6 fold during ultrasound application.^17^ We also targeted a median polydispersity index (PDI) of <0.1, to ensure monodispersity of each batch. We found that the PFP content in the reaction significantly affected the particle size, drug loading, monodispersity, and *in vitro* uncaging efficacy (Figs. S2, S3), and empirically determined that a 2 µl:1 mg ratio of PFP to polymer most reliably met our target size and PDI.

**Figure 1.**
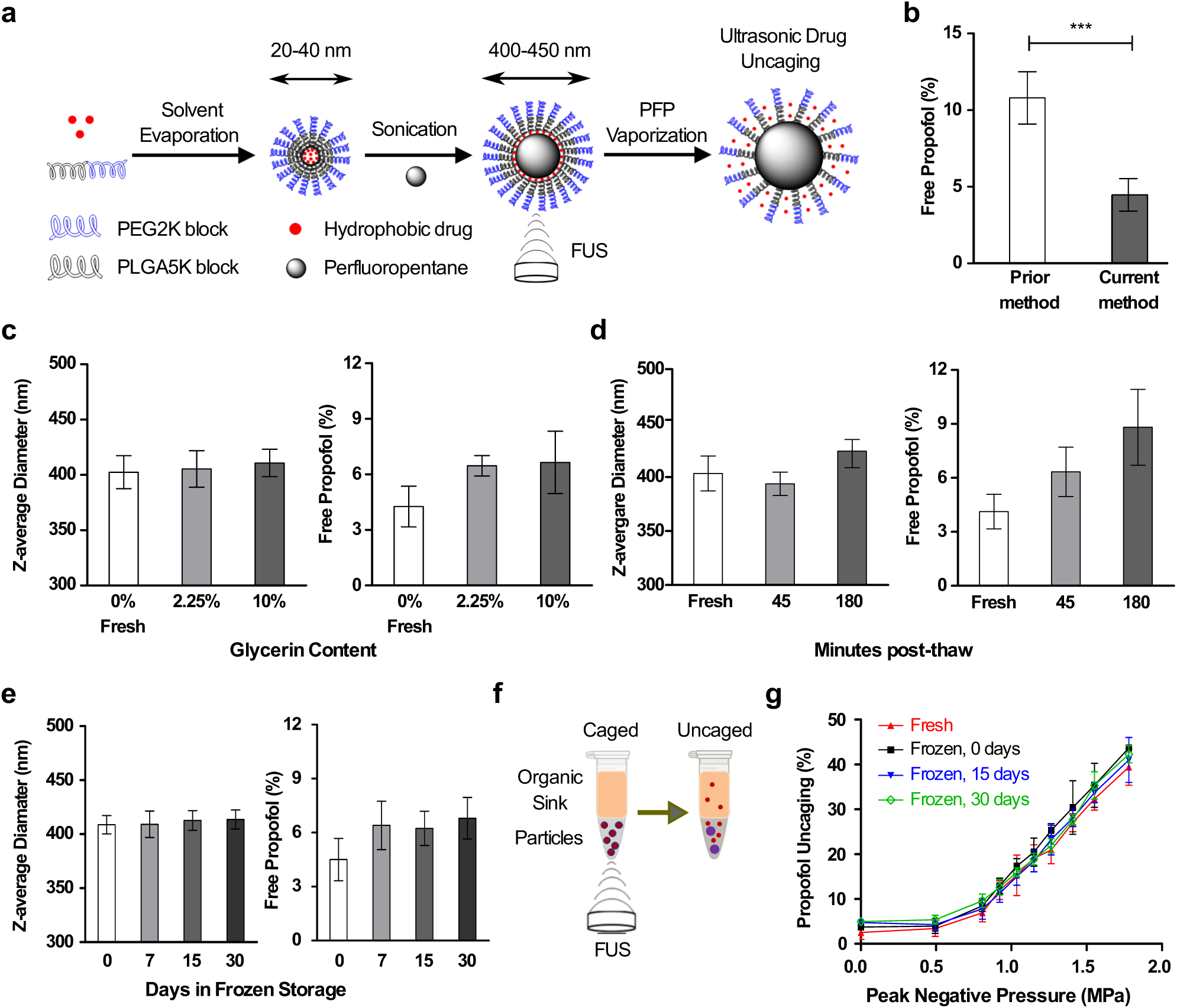
Optimized nanoparticle production for stability and efficacy *in vitro*. **(a)** Schematic of nanoparticle production and ultrasonic drug uncaging. **(b)** Free propofol content is reduced with the current optimized protocol versus the prior.^15^**(c-e)** Glycerin serves as a cryoprotectant to improve nanoparticle stability through frozen storage and thawing, compared to fresh (0, 0%) nanoparticle production. **(f)** Experimental schematic to assay ultrasonic drug uncaging efficacy *in vitro*. **(g)** Intact ultrasonic drug uncaging efficacy *in vitro* (650 kHz sonication, 60 × 50 ms pulses at 1 Hz pulse repetition frequency) for frozen & thawed nanoparticles compared to fresh. Mean +/- S.D. are presented for groups of N=3. ***: p < 0.001 by two-tailed t-test.

To generate the nanoparticles, the emulsifying polymer and drug were dissolved in tetrahydrofuran (THF), and then sterile phosphate-buffered saline (PBS) was added. The THF was evaporated to completion, leaving drug-loaded polymeric micelles (∼30 nm Z-averaged diameter) in saline suspension. Then, PFP was added and the mixture was sonicated in a bath sonicator until the PFP was visibly completely emulsified. Following three cycles of centrifugation and resuspension to remove free drug, polymer, and micelles, the nanoparticles were filtered twice through a membrane extruder to produce the final suspension. Notably, the shift from immersion sonication, as used previously,^15^ to bath sonication and membrane extrusion substantially reduced the free drug fraction (Fig. 1b) and minimized potential contamination due to exposure to the sonication probe. Dynamic light scattering confirmed that our current methods produced monodisperse peaks of nanoscale material (Fig. S1).

In prior perfluorocarbon nanoemulsion formulations, the particle size, free drug fraction, and polydispersity all increased substantially over the course of hours with incubation either on ice or at room temperature,^13,15^ and the particles were too unstable to permit a freeze-thaw cycle for long-term storage. To address this instability and significant practical limitation, we used cryoprotectants to enable frozen storage of the particles. The addition of minimal (2.25% w/v) glycerin to the particles had no substantial effect on the physicochemical characteristics and drug loading of the particles (Figs. 1c, S4a,b), yet allowed for improved particle stability in terms of physicochemical characteristics in the post-thaw time period (Figs. 1d, S4c,d) and permitted long-term frozen storage of the particles (Figs. 1e, S4e,f) with stability across multiple freeze-thaw cycles (Fig. S5). This formulation also showed low batch-to-batch variability with no change of physicochemical characteristics across varied particle concentrations (Fig. S4g-i, indexed by the encapsulated drug concentration). In our current protocol, there is a minimal slow increase of the free drug fraction during incubation, rising from ∼4% of the initial drug load when fresh to ∼8% after 3 hours post-thaw at room temperature (Fig. 1d).

To determine the *in vitro* efficacy of this novel formulation for ultrasonic drug uncaging, we loaded the particles into thin-walled plastic (PCR) tubes and then added a layer of organic solvent on top that is immiscible with and of lower density than water (Fig. 1f). Following focused sonication of the aqueous nanoparticle suspension, the organic layer was collected, and the UV spectra of this fraction was measured to indicate the amount of drug release. Indeed, there was robust FUS-induced drug release seen with a dose-response relationship with the applied *in situ* peak pressure, and no change of this efficacy between fresh and frozen/thawed nanoparticles, irrespective of the length of time that the particles were frozen (Fig. 1g). An inflection point for drug release versus sonication pressure was estimated as at 0.8 MPa for 650 kHz sonication (Figs. 1g, 3d) and at 1.2 MPa for 1.5 MHz sonication (Fig. 2d). Drug release also increased with sonication burst length, with minimal or no appreciable release below 10 ms and saturation of the effect above 50 ms.^18^

**Figure 2.**
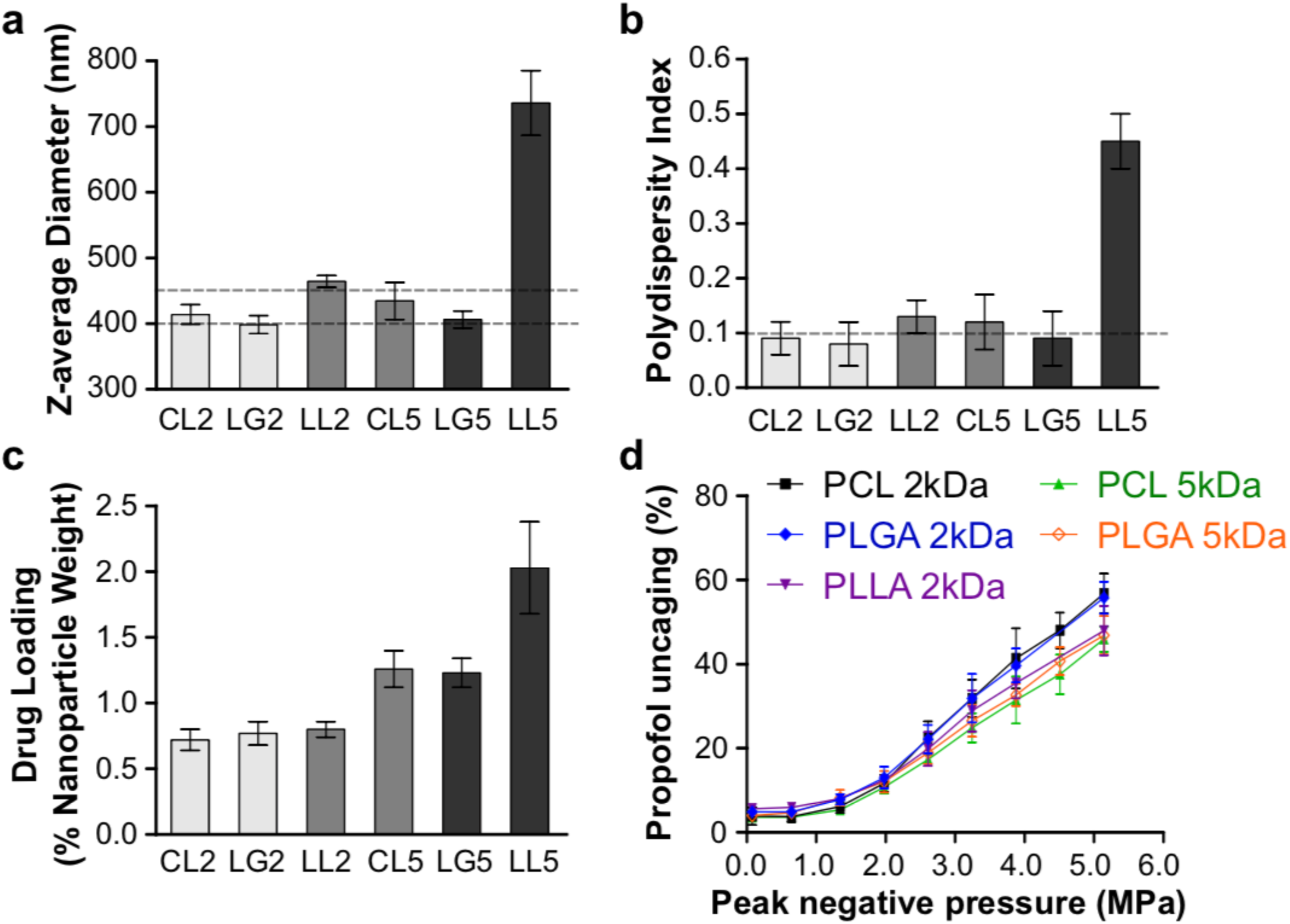
Optimization of polymer for drug-loaded perfluoropentane nanoemulsions. Diblock-copolymers were tested consisting of a hydrophilic block of PEG (2 kDa) and a choice of hydrophobic block among: PCL (2 kDa, CL2 or 5 kDa, CL5), PLGA (2 kDa, LG2 or 5 kDa, LG5), or PLLA (2 kDa, LL2 or 5 kDa, LL5) **(a)** Z-average diameter (dashed lines at the target values of 400-450 nm), **(b)** polydispersity index (dashed line at target value of 0.1), **(c)** propofol drug loading, and **(d)** ultrasonic propofol uncaging *in vitro* (1.5 MHz sonication, 60 × 50 ms pulses at s1 Hz pulse repetition frequency) was quantified. Mean +/- S.D. are presented for groups of N=3.

To evaluate the suitability of this production protocol for eventual clinical application, we assessed for endotoxin and bacterial contamination using a chromogenic LAL endotoxin assay and growth on LB agar plates, respectively. The endotoxin level measured 0.3 ± 0.05 EU per mg propofol, far lower than 5 EU per mg active pharmaceutical ingredient (API; propofol in this case, to be used at a target dose 1 mg propofol/kg body weight), which is considered to be acceptable for parenteral administration,^19^ indicating no significant endotoxin contamination. There was also no growth on LB agar plates of the nanoparticle formulation after 72 h at 37 °C, indicating the sterility achievable with this production protocol.

To determine the effect of the encapsulating diblock-copolymer on drug loading and release efficacy, we varied the hydrophobic block of the polymer between the common drug delivery polymers of poly-ε-caprolactone (PCL), poly-L-lactic acid (PLLA), and poly-lactic-co-glycolic acid (PLGA). The molecular weight of these blocks was varied between 2 kDa and 5 kDa. The hydrophilic block of poly-ethylene glycol (PEG; mol. wt. 2 kDa) was kept constant. PLLA particles, particularly with a block molecular weight of 5 kDa, showed increased size and polydispersity, and in many cases developed a precipitate during production (biasing the drug loading estimates), indicating that this polymer was not suitable for these applications (Fig. 2). This may be because the relatively long crystalline block of PLLA significantly reduces its solubility in the mixture of THF and PBS.^20^ PLLA was therefore removed from subsequent studies. There was minimal difference between PCL and PLGA in terms of the resultant particle physicochemical characteristics and drug loading, emphasizing the versatility of this approach for drug encapsulation. Larger hydrophobic blocks yielded greater drug loading (Fig. 2c), with approximately double the drug loading with 5 kDa hydrophobic block sizes compared to 2 kDa. There was minimal difference among the particles in terms of *in vitro* ultrasonic drug uncaging efficacy (Fig. 2d), with larger hydrophobic blocks trending towards minimally decreased percent uncaging. Given the substantially improved drug loading of 5 kDa vs. 2 kDa hydrophobic blocks relative to this minimally decreased percent uncaging, and the greater reported experience of safety and efficacy in clinical drug delivery applications with PLGA^21^ compared to PCL, we chose PEG(2 kDa)-PLGA(5 kDa) as the emulsifying polymer of the nanoemulsions for subsequent experiments.

### A Generalized Platform for Drug Delivery

To realize the promise of this system as a platform for targeted delivery of a wide variety of drugs, and to estimate the drug features that most enable encapsulation into polymeric perfluoropentane nanoemulsions, we varied the loaded drug across seven molecules: two vasoactive agents (calcium channel antagonists verapamil and nicardipine), three anesthetics (propofol, ketamine, and dexmedetomidine), and two chemotherapeutics (doxorubicin and cisplatin). There was minimal difference of the choice of loaded drug on the particle physicochemical properties (Figs. 3a,b). Instead, there was a strong positive relationship noted between the drug hydrophobicity (indicated by LogP, the oil:water partition coefficient) and the drug loading (Fig. 3c), with essentially no loading of the hydrophilic compound cisplatin. Interestingly, while there were minimal differences of *in vitro* ultrasonic drug uncaging efficacy across the different drugs (Fig. 3d), there was a reverse trend compared to drug loading in that doxorubicin (LogP=1.3) had marginally greater drug release versus applied pressure, compared to verapamil or nicardipine (LogP=3.8). These results establish the generalizability of this system for ultrasonic uncaging of hydrophobic drugs.

**Figure 3.**
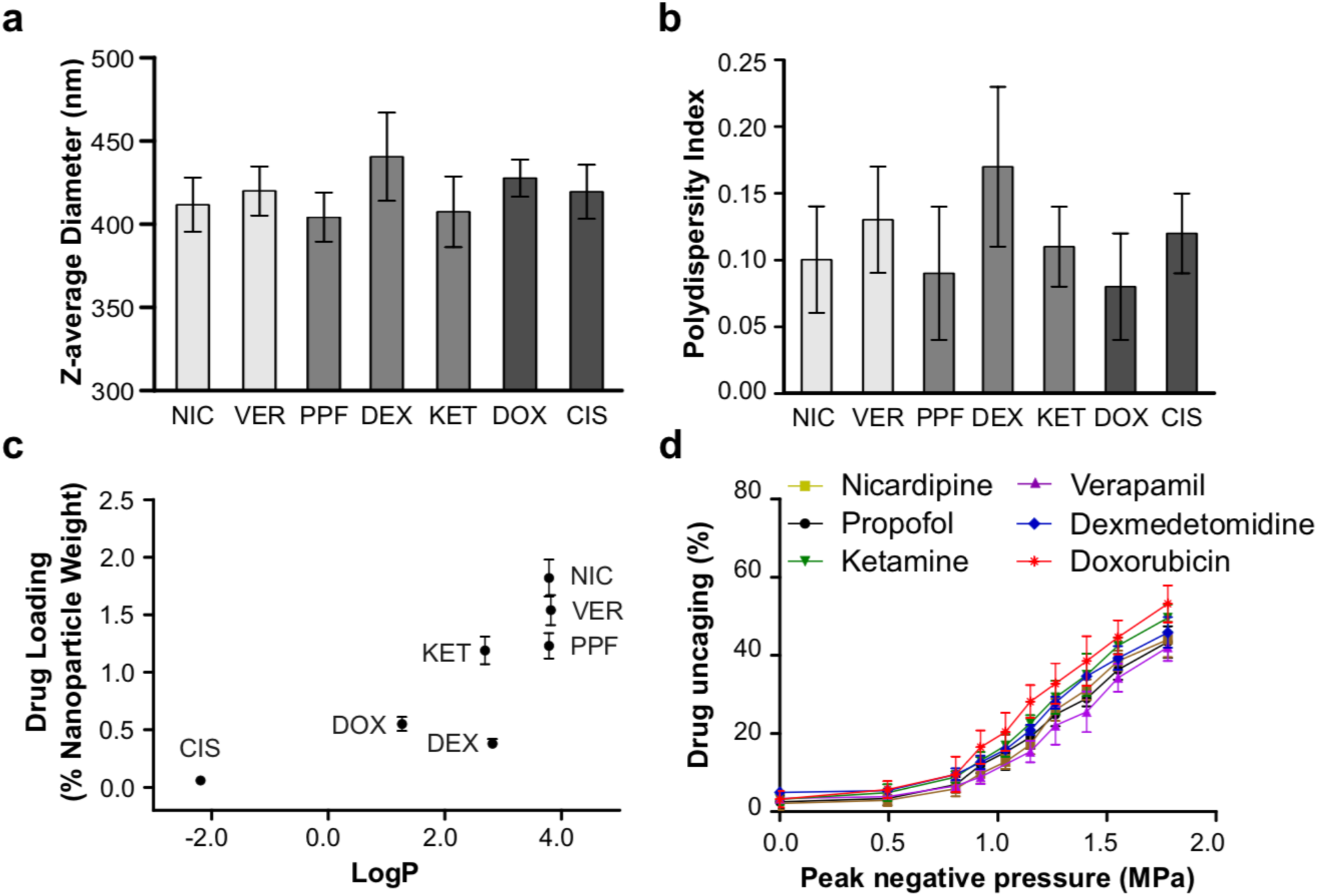
Polymeric perfluoropentane nanoemulsions are a platform for ultrasonic uncaging of hydrophobic drugs. **(a)** Z-average diameter, **(b)** polydispersity index, **(c)** drug loading, and **(d)** ultrasonic drug uncaging *in vitro* (650 kHz sonication, 60 × 50 ms pulses at 1 Hz pulse repetition frequency) of PEG(2 kDa)-PLGA(5 kDa) perfluoropentane nanoemulsions loaded with propofol (PPF), nicardipine (NIC), verapamil (VER), dexmedetomidine (DEX), ketamine (KET), doxorubicin (DOX), or cisplatin (CIS). Mean +/- S.D. are presented for groups of N=3. Cisplatin-loaded nanoparticles were not tested for *in vitro* uncaging due to low drug loading. In the bar charts, drugs of a similar class (vasodilators, anesthetics, chemotherapeutics) are grouped.

### *In Vivo* Nanoparticle Characterization

To determine the clearance kinetics, biodistribution, and biocompatibility of the particles in rats, in addition to the indicated drug, the particles were doped with a dye whose infrared fluorescence is quantitative in blood samples, and which clears from the blood pool within ∼5 min in its unincorporated free form. For this analysis, we chose among the drugs with substantial loading (Fig. 2), drugs with high (nicardipine), intermediate (propofol), and low (doxorubicin) drug loading. To assess particle clearance kinetics, the fluorescence was quantified for whole blood and plasma samples collected at several time points over hours. The difference between the whole-blood and plasma sample fluorescence indicated the nanoparticle blood concentration. The plasma fluorescence indicated the rate of generation of drug-loaded micelles as the volatile PFP diffuses out of the nanoparticle core, as well as the potential elution of free dye. There was no substantial effect of the choice of encapsulated drug on particle kinetics or biodistribution (Fig. 4). Independent of the particular loaded drug, the particle blood pool concentration followed a dual exponential clearance profile, with half-lives of 10-12 min and 77-97 min for each phase (Fig. S6; Tables S1-2). Based on this profile, a bolus plus infusion protocol was determined to yield a steady blood particle concentration to enable prolonged usage. Indeed, with this bolus plus infusion protocol, a steady blood pool particle concentration was seen for over 40 min, with a similar elimination profile to bolus alone following the halt of infusion (Fig. 4b). Independent of the particular loaded drug, the particles showed uptake at 24 h primarily in the liver, followed by spleen and lung, with minimal uptake in kidney and heart, and notably no binding to the brain (Fig. 4e-f). In our experiments, >100 rats have received the current formulation of these particles, with some receiving up to nine doses over several weeks, and none has shown visible evidence of toxicity due to particle administration or uncaging. Indeed, no negative change was seen in animal body weight across two weeks of multiple nanoparticle administrations (Fig. 4g). The brains of animals that underwent uncaging in the brain following nanoparticle administration and uncaging sonication showed no evidence of acute injury or blood-brain barrier disruption by either histology or contrast-enhanced MRI.^21^ These results indicate that, independent of the choice of loaded drugs, these nanoparticles and their uncaging are well tolerated and have clinically practical clearance kinetics to enable acute ultrasonic drug uncaging therapies.

**Figure 4.**
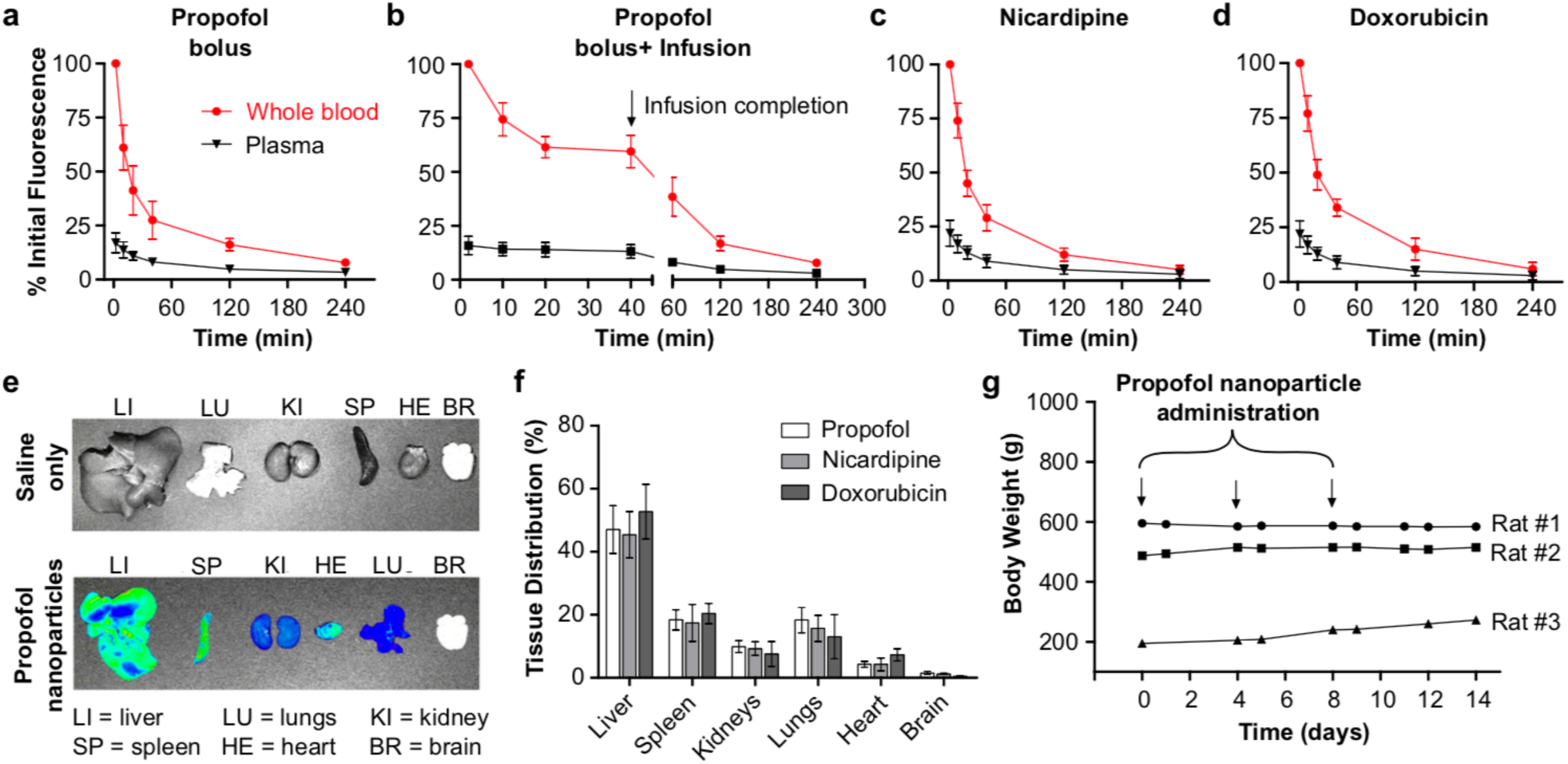
Particle clearance kinetics, biodistribution, and tolerance. Particle kinetics after intravenous administration of **(a)** propofol-loaded nanoparticles as a bolus (1 mg/kg encapsulated propofol), **(b)** propofol-loaded nanoparticles as an i.v. infusion (bolus of 1 mg/kg + infusion of 1.5 mg/kg/hr encapsulated propofol), **(c)** nicardipine-loaded nanoparticles as a bolus (1 mg/kg encapsulated nicardipine), and **(d)** doxorubicin-loaded nanoparticles as a bolus (1 mg/kg encapsulated doxorubicin). **(e)** Sample images of IR dye fluorescence in organs harvested 24 h after saline (top) and nanoparticle (bottom) administration to rats. **(f)** Tissue distribution of propofol, nicardipine, or doxorubicin-loaded nanoparticles 24 h after i.v. bolus (1 mg/kg encapsulated drug). **(g)** Body weight of rats administered 3 boluses of propofol-loaded nanoparticles over 8 days. On Day 0, Rat #1: 15 weeks old and body weight 595 g; Rat #2: 11 weeks old and body weight 457 g; Rat#3: 5 weeks old and body weight 195 g. Mean +/- S.D. are presented for groups of N=3.

### Quantification of *In Vivo* Drug Pharmacokinetics with Ultrasonic Uncaging

Given the favorable kinetics of particle clearance and biodistribution that are independent of the loaded drug, we then assessed the *in vivo* pharmacokinetics of ultrasound-induced drug release for this system. We first note that no prior demonstration of perfluorocarbon nanoemulsion uncaging, including our own work, demonstrated that this uncaging yields a restricted pharmacokinetics or pharmacodistribution of drug release. For this current assay, the plasma concentration of drugs was determined in blood samples taken before and after ultrasound application. In a first scenario, ultrasound was applied to the lower abdominal aorta and blood was sampled from the downstream femoral vein (Fig. 5a-d). In a second scenario, ultrasound was applied to the brain (frontal cortex), similar to our previous implementation of this uncaging,^15^ and blood was sampled from the downstream ipsilateral internal jugular vein (Fig. 5e-f). In each scenario, for each drug, while before sonication there was no plasma drug concentration significantly above the detectable limits for this assay, following ultrasound (240 × 50 ms pulses, 1 Hz pulse repetition frequency, 1.5 MPa estimated peak *in situ* pressure for abdominal aorta or 1.2 MPa estimated peak *in situ* pressure for frontal cortex) a sharp rise in plasma drug concentration was noted in the downstream vein, with a dual-exponential decay profile of the plasma drug concentration. Notably, the pharmacokinetics of the released drug and the estimated dual-exponential decay half-lives (Table S3) were similar whether uncaging was completed in the largest artery of the body (the aorta, Fig. 5a-d) or directly in organ parenchyma where the uncaging would occur in the limited blood volume of the capillary bed (Fig. 5e-f). This indicates that the drug in either case is mostly extracted during a first pass of perfusion, independent of the loaded drug or the sonication site, underlining the specificity and versatility of this drug delivery technique and confirming that ultrasonic drug uncaging from perfluorcarbon nanoemulsions yields pharmacokinetics that are limited in time by ultrasound application.

**Figure 5.**
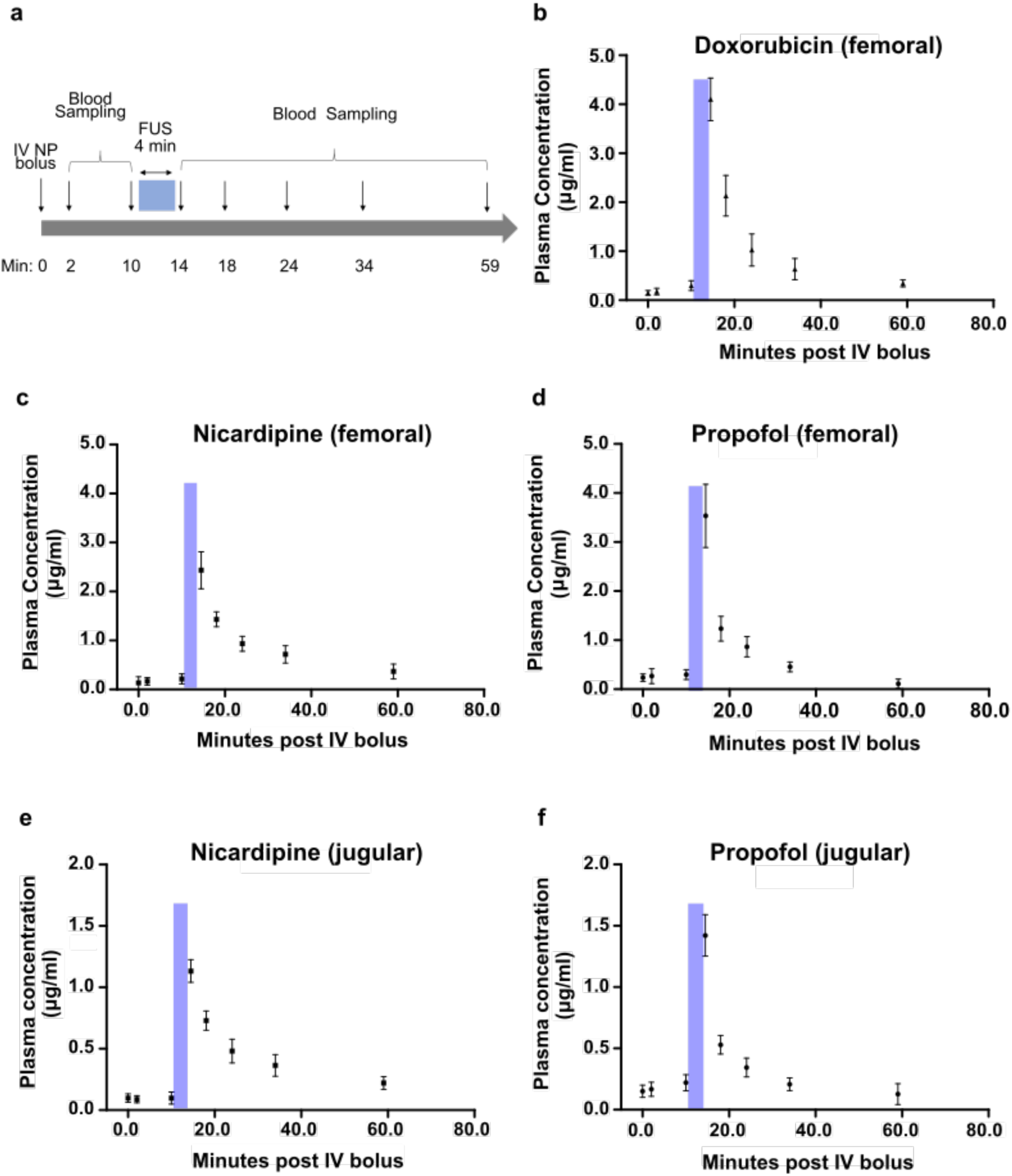
Drug pharmacokinetics following ultrasonic drug uncaging. **(a)** Experimental schematic of blood sampling before and after FUS (240 × 50 ms pulses, 1 Hz pulse repetition frequency, estimated *in situ* peak pressure of 1.5 MPa for femoral sampling, 1.2 MPa for jugular sampling). **(b-d)** Plasma drug profile following sonication of the lower abdominal aorta and blood sampling of the left femoral vein. **(e-f)** Plasma drug profile following sonication of the brain frontal cortex and blood sampling of the ipsilateral internal jugular vein. Blue bars indicate FUS timing. Samples at t=0 min were taken immediately prior to nanoparticle infusion via a tail vein and represent the assay sensitivity limits. Presented are mean +/- S.D. for groups of N=3.

### Efficacious and Localized *In Vivo* Ultrasonic Drug Uncaging

Next, we sought to confirm that these particles retained high therapeutic efficacy *in vivo*. In our previous work, we verified that propofol-loaded perfluorocarbon nanoemulsions could induce drug uncaging efficacious enough to silence chemically-induced whole-brain seizures.^15^ Notably no work has demonstrated a spatially restricted drug effect with uncaging from perfuorocarbon nanoemulsions, neither that of our own nor of others.

Additionally, given that drug uncaging in this technique occurs intravascularly, we note that the broadest pharmacodistribution would occur with uncaging directed to an artery that subtends a large region of the body, especially for a drug that mainly acts on the vessel itself, instead of in a target organ. Notably vasodilating agents, such as nicardipine and verapamil, are used clinically to relieve arterial spasm as seen with cerebral vasospasm and other conditions,^22^ by relaxing the smooth muscle of the vessel wall. For example, nicardipine has been shown to relax the wall of the aorta and increase its distensibility in humans.^23^ While effective, these agents have undesirable side effects of generalized hypotension when given systemically, due to decreasing the systemic vascular resistance by action beyond the target vessel. This hypotension can result in end organ infarction in severe cases. In order to minimize this effect, the vasodilator must be infused via an invasive intra-arterial catheter placed within the target vessel or immediately upstream. For ultrasonic vasodilator uncaging to achieve similar effects, the vasodilator must bind the arterial smooth muscle immediately after ultrasound-induced release from the nanoparticles, given that arterial velocities are generally on the order of 0.3-0.5 m/s.^24^ Therefore, to stress-test whether ultrasonic drug uncaging from perfluorocarbon nanoemulsions could achieve a local and substantial drug effect, we assessed whether ultrasonic nicardipine uncaging could yield a visible change in vessel wall compliance of the rat abdominal aorta (∼7 cm in length, ∼1 mm luminal diameter), the highest flow vessel of the body. With ultrasonic nicardipine uncaging in the aorta upstream of an ultrasound imaging probe, a substantial difference in systolic and diastolic aortic diameters was noted compared to the pre-uncaging baseline (Fig. 6; Videos S1-2). Confirming its specificity, this effect was not seen with ultrasound alone, with nanoparticle administration alone, with ultrasound applied to blank nanoparticles, or with nicardipine uncaging applied downstream of the imaging probe (‘Position 2’; Fig. 6a,c). In fact, compared with a systemic bolus of free nicardipine that is matched in terms of the total nicardipine dose, ultrasonically uncaged nicardipine had a more potent effect on the vessel wall distensibility (Fig. 6c), even though it is likely that only a minority of the nanoparticles were exposed to the sonication field. Additionally, systemic nicardipine administration increased the aortic blood velocity by 41% on average, due to a decrease in systemic vascular resistance with peripheral nicardipine action (Fig. S7). Similarly, uncaging of nicardipine nanoparticles in the distal aorta (i.e. ‘Position 2’) increased the blood flow velocity in the aorta (Fig. S6), likely by relaxing the arterial/arteriolar smooth muscle in the lower limbs and therefore decreasing the vascular resistance seen by the aortic blood flow. However, importantly, this effect was not seen with ultrasonic nicardipine uncaging in the proximal aorta (‘Position 1’), confirming that ultrasonic drug uncaging is limited to effects in the immediate region of sonication, given the length of the rat aorta (∼7 cm) and the rapid speed of rat aortic flows (∼0.3-0.5 m/s; Fig. S7). This confirms that ultrasonic drug uncaging yields a volume of distribution of the drug that is effectively confined to the sonicated region, which in turn results in an effective amplification of the local drug effect. Furthermore, these results demonstrate that this localized drug-receptor binding can occur even in the presence of rapid aortic flows. Notably, the majority of this effect occurred with the first minute of sonication (Fig. 6d), confirming the rapid temporal kinetics of the bioeffects of ultrasonic drug uncaging (Fig. 5).

**Figure 6.**
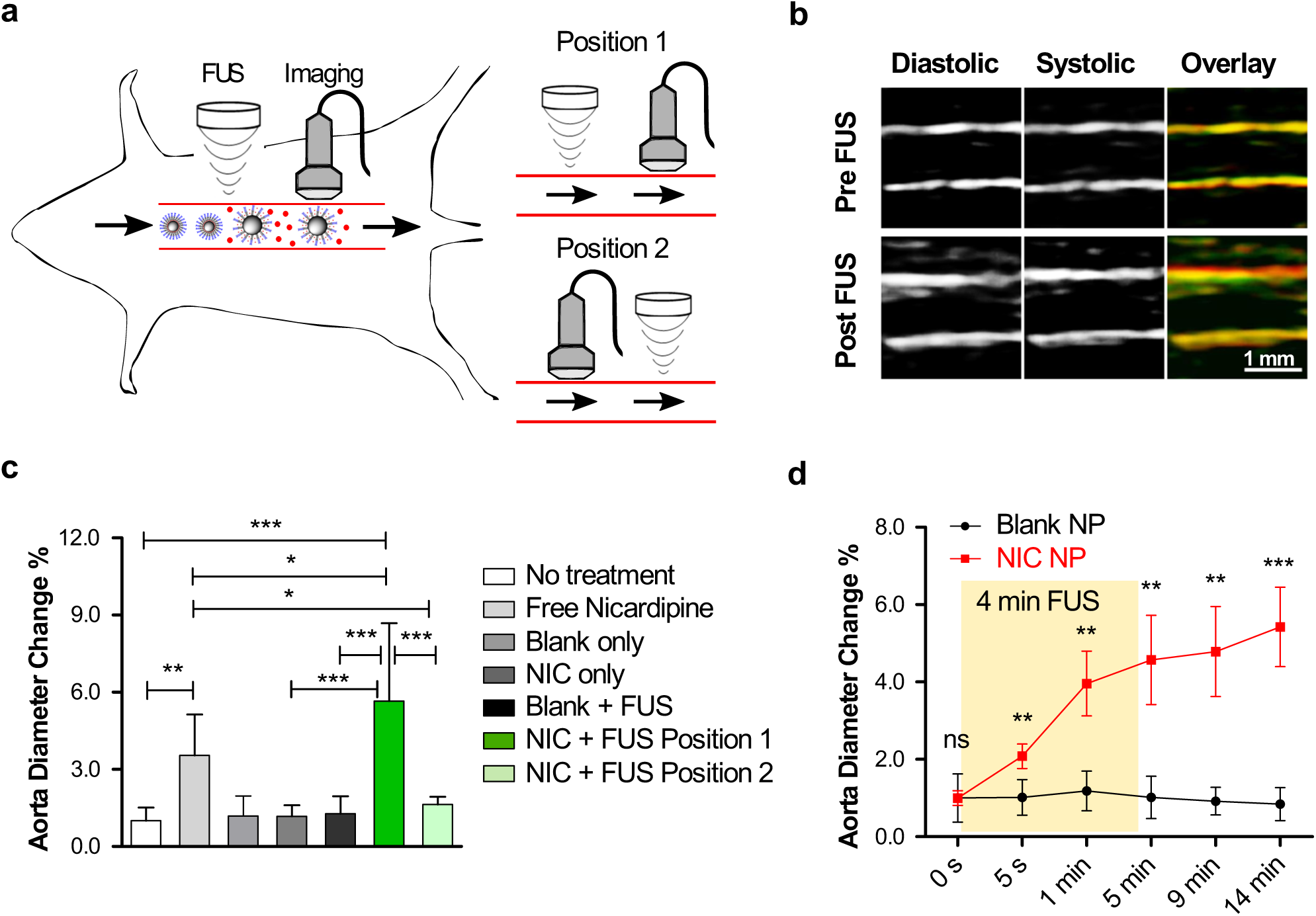
Ultrasonic nicardipine uncaging increases local vessel compliance. **(a)** Experimental schematic to test if ultrasonic nicardipine uncaging from nanoparticles increases rat aortic wall compliance. Uncaging is applied to the aorta either upstream (Position 1) or downstream (Position 2) of imaging. **(b)** Ultrasound images of the rat abdominal aorta during systole and diastole, before and after ultrasonic nicardipine uncaging (650 kHz, 240 × 50 ms pulses at 1 Hz pulse repetition frequency, 1.5 MPa est. peak *in situ* pressure); green = diastolic, red = systolic, yellow = green/red overlap. Averaged rat abdominal aortic diameter at **(c)** 14 min after uncaging or **(d)** across time, normalized by the initial (0 s) values. FUS = focused ultrasound application, NIC = nicardipine-loaded nanoparticles. Free nicardipine and NIC administered to total drug dose of 134 µg/kg i.v. Mean +/- S.D. are presented for groups N=5-6 (**c, d**). ns: not significant, **: p < 0.01; ***: p < 0.001 by ANOVA and Tukey post-hoc tests (**c,** F(6,30)=28.49) or by two-tailed Student’s t-tests between the nicardipine-loaded nanoparticles and the corresponding negative conditions (**d**).

## Discussion

We have shown that polymeric perfluorocarbon nanoemulsions are a versatile platform for ultrasonic drug uncaging, with a ready path for clinical translation. We have described scalable production methods that hew to clinical standards and produce particles that are stable for both long-term frozen storage and for hours of use after thawing (Figs. 1, S1-5). We have confirmed that longer hydrophobic blocks of the emulsifying polymer yield greater drug loading, with minimal effect of the specific choice of polymer on drug uncaging efficacy (Fig. 2). We further explicitly demonstrate the ability of this technology to encapsulate and selectively uncage drugs that span a range of hydrophobicity, drug classes, and receptor binding profiles (Fig. 3). Indeed, while the drug loading increases with drug hydrophobicity, all other features of the particles, including their drug release efficacy, clearance kinetics, and biodistribution are relatively independent of the particular drug that is encapsulated (Figs. 3, 4, S6; Tables S1-2). Notably, *in vivo* ultrasonic drug uncaging with these particles produces a restricted pharmacokinetics (Figs. 5; Table S3) and a localized pharmacodistribution following intravascular release of the drug cargo (Figs. 6, S7, Videos S1-2). Combined with our prior results showing efficacy of this system for application in the brain,^15^ these results indicate that ultrasonic drug uncaging with perfluorocarbon nanoemulsions provides a noninvasive and wireless analogue to catheter-based intravascular infusion of hydrophobic drugs.

There are numerous potential applications for ultrasonic drug uncaging. In the brain, focal uncaging of neuromodulatory agents^25^ could allow pharmacological adjuncts to psychiatric therapy sessions that are tailored to the particular neural circuit pathophysiology for a given patient. This technology could also allow pharmacological mapping of neural circuits to better target more permanent interventions such as surgical resection, ablation, or deep-brain stimulation. Additionally, current therapy for vessel spasm disorders, such as the cerebral vasospasm that unfortunately accompanies many cases of subarachnoid hemorrhage,^22^ is difficult given that the agents that best relieve the spasm also act systemically as potent anti-hypertensives. The noninvasive local relaxation of the walls of the affected vessels, as modeled in Fig. 6, would be enormously beneficial as a noninvasive alternative to current catheter-based intra-arterial vasodilator infusions, especially as this nanotechnology achieves local vasodilatation without concomitant loss of systemic vascular resistance (Fig. S7) and therefore systemic hypotension. Finally, many chemotherapeutics are known to be effective for treatment of a given tumor cell type, yet cannot be administered in effective doses systemically due to intolerable side effects in the rest of the body. Ultrasonic chemotherapeutic uncaging within the tumor and its immediate margin would therefore be of great utility.

Future work with this technology will move ultrasonic drug uncaging to clinical practice, first by validating this approach in large animal models, and then by beginning first-in-human trials to establish the safety and the efficacy of drug uncaging of this technique. Importantly, the constituent components of these nanoparticles – namely the drugs under consideration, PEG, PLGA,^21^ and PFP^16^ – have each been used in clinical populations with excellent safety profiles, lowering the barrier to translation for these nanoparticles. Additional future work will focus on expanding this technology to include encapsulation of hydrophilic small molecules, as well as larger macromolecules like peptides, antibody fragments, and nucleic acids. Given their potential for clinical translation, their ability to uncage a variety of important drugs, and the potent local pharmacological bioeffects they can induce in the brain^15,21^ and in the body, polymeric perfluoropentane nanoemulsions are poised to have enormous impact both for clinical care as well as our scientific understanding of how pharmaceuticals mediate their bioeffects.

## Conclusions

We have developed polymeric perfluoropentane nanoemulsions as a versatile platform for targeted drug delivery that is primed for clinical translation. These nanoparticles are produced with clinically compatible production methods and have stability, drug loading, and drug release efficacy that is suitable for clinical application. This system efficaciously encapsulates and ultrasonically uncages a wide variety of drugs, with drug loading increasing with hydrophobicity, and with all other features of the particles being independent of the particular encapsulated drug. Further, the *in vivo* pharmacokinetics and pharmacodistribution of drug release with this technology can be defined according to when and where ultrasound is applied. In this manner, polymeric perfluorocarbon nanoemulsions can be used for wireless intravascular hydrophobic drug infusions, as noninvasive analogues of current catheter-based therapies.

## MATERIALS AND METHODS

### Materials

Di-block copolymers are made up of a hydrophilic block of polyethylene glycol (PEG; mol. wt. 2 kDa) and a hydrophobic block of one of: poly(lactic-co-glycolic acid) (PLGA), poly(L-lactic acid) (PLLA), or poly(ε-caprolactone) (PCL). Two molecular weights of hydrophobic block chains were used: 2 kDa and 5 kDa. The example of nomenclatures for di-block copolymer is polyethylene glycol 2 kDa - poly(lactic-co-glycolic acid) = PEG (2 kDa)-PLGA (5 kDa). All di-block copolymers were purchased from Akina (West Lafayette, IN, USA). Propofol, nicardipine hydrochloride, verapamil hydrochloride, sodium sulfate and sodium hydroxide were purchased from Alfa Aesar (Haverhill, MA, USA). Doxorubicin hydrochloride was purchased from LC laboratories (Woburn, MA, USA). Cisplatin, dexmedetomidine, Luria Broth (LB) powder, and LB agar powder were purchased from Sigma-Aldrich (St Louis, MO, USA). Ketamine hydrochloride injectable solution is a controlled substance and was purchased via Stanford University Environmental Health & Safety. Tetrahydrofuran (THF), methanol, ethyl acetate, chloroform, and hexane were obtained from Sigma-Aldrich (St Louis, MO, USA). n-Perfluoropentane (PFP) was purchased from FluoroMed (Round Rock, TX, USA). A hydrophobic IRDye® 800RS infrared dye was purchased from LI-COR Biotechnology (Lincoln, NE, USA).

### Production of Drug-Loaded Polymeric Perfluoropentane Nanoemulsions

The production of polymeric perfluoropentane (PFP) nanoemulsions was similar for all the tested drugs and amphiphilic di-block copolymers. Briefly, 150 mg of di-block copolymer and 15 mg of drug with hydrochloride removed (Method S1 in Supporting Information) were weighed into a 20 ml glass beaker and 10 ml THF was added to dissolve the polymer and drug. Then, 10 ml phosphate buffer saline (PBS) was added dropwise to the organic solution over 5 min. The THF was fully evaporated by placing the mixture overnight in atmosphere and then in vacuum for 1 h. Then, 300 µl cold PFP (volume fraction = 3% relative to micelle solution) was added to the micelle solution, followed by 5 min sonication in a 40 kHz Bransonic M1800H bath sonicator (Thermo Scientific; Waltham, MA, USA) which was pre-filled with iced water. The resulting nanoparticles were washed three times and collected by a total of 10 min centrifugation 4 °C (2000 RCF). Finally, the nanoemulsion suspension was extruded using an Avestin Liposofast LF-50 extruder (Ottawa, ON, Canada) equipped with compressed nitrogen (40-100 psi) and loaded with a Nuclepore Track-Etch polycarbonate membrane of 0.6 µm pores (GE Healthcare, Chicago, IL, USA). The extruded nanoemulsion suspension was either used fresh or mixed with glycerin (2.25%, w/v) and frozen immediately and stored at −80 °C.

### Physicochemical Characterization of Drug-Polymeric Perfluoropentane Nanoemulsions

The Z-average diameter, polydispersity index (PDI) and zeta potential of the drug-loaded phase-change nanoemulsions were measured with a Malvern Zetasizer Nano ZS90 (Malvern, United Kingdom). The drug loading in the nanoemulsion was quantified with either UV absorption or fluorescence. The details can be found in the Method S2-3 of Supporting Information. Endotoxin concentration of the propofol-loaded nanoemulsion was assayed with ToxinSensor^™^ LAL endotoxin kit (GenScript. NJ, USA) following the provided protocol (Protocol: L00350; Supporting Information S4). According to the US Food and Drug Administration, ≤5 EU/mg drug is considered an acceptable endotoxin concentration for propofol intended to be delivered to a dose of 1 mg/kg.^19^ The sterility of the propofol-loaded nanoemulsion was evaluated by plating on LB agar plates and assessing colony growth at 72 h of incubation at 37 °C.

### Nanoemulsion Stability at Varied Temperatures

Propofol-load nanoemulsions were used to assess the particle stability at different temperatures. Z-average size, polydispersity index and free propofol content in the nanoemulsion were evaluated after frozen storage at −80 °C and at 0 °C, after thaw. The nanoemulsion was assessed after 7, 15 and 30 days in storage at −80 °C. After thaw from −80 °C, the above-mentioned parameters of nanoemulsion were assessed at 45 min and 3 hrs after thaw. We also studied the effect of the concentration of nanoemulsion (as indexed by drug concentration) on particle stability during storage. The initial concentration of propofol in the nanoemulsion was selected as either 0.5, 1, or 3 mg/ml by adjusting the resuspension PBS volume in nanoemulsion production. Finally, we assessed whether repeated freeze-thaw treatment impacts the integrity of nanoemulsion. The nanoemulsion was thawed as described above and then frozen shortly after sampling. Five freeze-thaw cycles were performed consecutively.

### *In Vitro* Assay of Ultrasonic Drug Uncaging

The effect of polymer composition and drug partition coefficient (LogP) on *in vitro* drug uncaging from nanoemulsions was studied. Propofol was used as a model drug to study the effect of the varying hydrophobic polymer blocks. A 50 µl nanoemulsion suspension (1 mg/ml drug equivalent) was added to a Fisherbrand™ 0.2 ml PCR tube (Fisher Scientific. NJ, USA). A 150 µl organic solvent of density less than water was added atop the nanoemulsion suspension. The exact solvent used varied depending on the drug being tested: hexane was used to extract propofol and ketamine; ethyl acetate was used for nicardipine, verapamil, dexmedetomidine, and doxorubicin. The PCR tube was placed in a custom holder and coupled using degassed water to a focused ultrasound (FUS) transducer (Image Guided Therapy, Pessac, France) at room temperature, so that the FUS focus was contained within the nanoemulsion suspension layer (Fig. 1f). The nanoemulsions were sonicated with FUS for 60 s total, with varying peak negative pressure, using cycles of 50 ms ultrasound on and 950 ms off, i.e. pulse repetition frequency of 1 Hz. The center frequency of the transducer was 1.5 MHz or 650 KHz. Following FUS, 100 µl of the organic solution was collected without disturbing the aqueous layer. The amount of the uncaged drug was quantified by measuring its UV or fluorescence and comparing to a standard curve of the drug prepared in varying concentrations in the same organic solvent. PEG (2 kDa)-PLGA (5 kDa) was used to create all nanoemulsions for the analysis of how the drug LogP affects nanoemulsion characteristics. The experimental setup and procedure were otherwise similar.

### Pharmacokinetics and Biodistribution of Drug-Loaded Polymeric PFP Nanoemulsions

All animal experiments were carried out in accordance with the Stanford IACUC. Long-Evans rats with body weight 180-200 g (Charles River Laboratories, Wilmington, MA, USA) were used in all *in vivo* studies. Drug-loaded PFP/PEG (2 kDa)-PLGA (5 kDa) nanoemulsions were doped with a hydrophobic near infrared fluorescent dye - IR800, during nanoemulsion production. Propofol, nicardipine, and doxorubicin-loaded nanoemulsions were used to test *in vivo* blood-pool nanoparticle kinetics and systemic biodistribution.

To produce dye-doped nanoemulsions, 1 mg IR800 dye was added to the drug and polymer THF solution, and the rest of the nanoemulsion production protocol was unchanged. For the experiments, a nanoemulsion bolus (equivalent to 1 mg/kg of drug) was administered intravenously via a 24 g × 3/4” catheter to rat tail vein in a total volume of ∼0.4-0.5 ml (N=3). Blood samples were collected via left and right submandibular veins at 2 min 10 min, 20 min, 40 min, 2 h and 4 h, alternating sides for each sampling. The blood was split into two volumes. Whole blood sample fluorescence was assessed using a Lago (Spectral Instruments Imaging; Tucson, AZ, USA) imaging system (excitation/emission = 770/810 nm) and quantification was completed using regions of interest (ROIs) of the same size across samples, drawn to be within the capillary tube. The second volume of each sample was centrifugated in a microcentrifuge for a total of 10 min at 10,000 g at 4 °C. The plasma fraction from these samples was then collected and their fluorescence was quantified similar to that of whole-blood samples. The nanoemulsion concentration in the whole blood and plasma were fitted with a two-compartment kinetic model. The clearance kinetics of dye-doped propofol-loaded nanoemulsions administered as a bolus (equivalent to 1 mg/kg of propofol) followed by an immediate infusion (equivalent to 1.5 mg/kg/hr of propofol) was also quantified. Blood was collected and quantified as described above.

For systemic biodistribution, the same dye-doped nanoemulsions (propofol, nicardipine, or doxorubicin-loaded) were administered intravenously as a bolus to Long-Evans rats (N=3). The rats were sacrificed at 24 h post administration to harvest major organs: heart, liver, lungs, kidneys, spleen, and brain. These organs were imaged for IR800 fluorescence (Ex/Em=770/810 nm) using a Lago imaging system and quantified using regions of interest (ROI) of the same size, drawn to be within the image of each organ. The distribution of the nanoemulsion among the organs was calculated by dividing the ROI fluorescence of each tissue by the sum of ROI fluorescence values of all organs.

### *In Vivo* Uncaging of Nicardipine in Abdominal Aorta

Long-Evans rats (N= 5-6) were positioned in a supine position and the hair overlying the abdomen was removed with electric clippers. The cranial portion of the abdominal aorta was imaged to determine the location of the sonication target. Once confirmed, the FUS probe was positioned overlying the target abdominal aorta. Separately, the imaging transducer was positioned over the distal aorta just proximal to the aortic bifurcation (corresponding to Position #1; Figure 6a). B-mode and power Doppler imaging were performed with a Siemens Acuson S2000 scanner with a Siemens 15L4 transducer using a transmit frequency of 10 MHz for B-mode and 7.5 MHz for power Doppler. For experiments where the probes were reversed (i.e. Position #2; Figure 6a), the imaging transducer was positioned in the upper abdominal aorta, and the FUS probe was positioned to target the distal aorta. Once the imaging probe and FUS sonication probe were placed, a bolus (∼0.3-0.5 ml) of nicardipine-loaded (equivalent to 134 µg/kg nicardipine) or blank nanoemulsions was slowly given through a tail vein catheter. Then, the FUS sonication was initiated at an estimated peak in situ pressure of 1.5 MPa, with pulsed sonication delivered as 50 ms on and 950 ms off (i.e. 1 Hz pulse repetition frequency) for 240 pulses (i.e. 4 min in total). The center frequency of the transducer was 650 KHz (Image Guided Therapy, Pessac, France). At this frequency, with 0.54 dB/cm*MHz of soft tissue attenuation, and typically 7 mm of soft tissue overlying the aorta, we estimate a net attenuation of −0.25 dB, or 3%, of pressure. The aortic distensibility was measured by calculating the percent change in the maximum diameter of the inner aortic wall between the cardiac systole and diastole phases, as shown in the following equation: Aorta distension %=[(diastolic diameter-systolic diameter)_post treatment_/(diastolic diameter-systolic diameter)_pre treatment_ - 1]×100%. The treatment in the equation denotes the i.v. injection of nanoparticles or free nicardipine followed by FUS. The measurements were made by two independent observers who were blinded to the experimental condition, and the averaged values were used for analysis.

### Quantification Plasma Concentration of Ultrasonically Uncaged Drug

The left femoral vein and tail vein of Long-Evans rats (N=3) was cannulated and then the rat was placed in a supine position and the hair overlying the abdomen was depilated. The ultrasound imaging probe and FUS transducer were placed at “Position 2” (see previous section). Then, 1 mg/kg drug equivalent nanoemulsion (propofol, doxorubicin, or nicardipine-loaded nanoemulsion) was administered via the tail vein. A 0.25 ml blood was sampled via the cannulated femoral vein at 2 and 10 mins after intravenous injection. The FUS was then delivered to the lower abdominal aorta from 10 to 14 mins after bolus injection. The FUS condition were same as that in the prior section. A 0.25 ml blood was then sampled at 14.5, 18, 24, 34, and 59 mins after intravenous injection, corresponding to 0.5, 4, 10, 20, and 45 mins after FUS.

A 0.1 ml plasma sample was obtained by a total of 10 min centrifugation at 5, 000 g at 4 °C for each blood sample. The drug was then extracted with 0.2 ml organic solvent, which is hexane for propofol, and ethyl acetate for nicardipine and doxorubicin. The propofol content in hexane was quantified with fluorescence (Ex/Em= 276/292 nm). For nicardipine, the UV absorbance of extracting ethyl acetate was measured at 348 nm. For doxorubicin, was measured at 494/595 nm. The drug concentration was calculated using a pre-determined standard curve of drug fluorescence or UV absorbance with respect to drug concentration in the corresponding solvent.

A similar *in vivo* drug uncaging of propofol and nicardipine was also performed by sonication on the brain (i.e. frontal cortex), followed by blood sampling from a cannulated left internal jugular vein. The animals were placed in a stereotactic frame (Image Guided Therapy, Pessac, France) coupled to the FUS system, and immobilized with two ear bars and a bite bar. The FUS transducer was aligned to 7 mm anterior to the interaural line, 2 mm lateral of midline. The blood collection timeline and sonication protocol were similar. The *in situ* pressure was approximately 1.2 MPa accounting for the attenuation by rat skull.

### Statistical Analysis

GraphPad Prism 5 (GraphPad Software, La Jolla, CA, USA) was used for statistical analysis. Comparisons between two groups was performed by two-tailed Student’s t test, and that among multiple groups by one-way analysis of variance (ANOVA) with *Tukey’s post-hoc* test. A p-value of 0.05 or less was considered statistically different and the *p*-values were categorized as *:*p*<0.05, **:*p*<0.01, ***:*p*<0.001, and ****:*p*<0.0001.

## ASSOCIATED CONTENT

### Supporting Information

i. Supplementary methods of dynamic light scattering, drug loading quantification, endotoxin assay, and LB agar plate test.
ii. Figure S1: Typical dynamic light scattering spectrum of propofol-loaded PFP polymeric nanoemulsion;
iii. Figure S2: Physicochemical properties of polymeric nanoemulsions with varied volume ratios of PFP to nanoparticle;
iv. Figure S3: Effect of particle size on in vitro uncaging efficacy of propofol-loaded nanoemulsions;
v. Figured S4-S5: Physicochemical property changes of propofol-loaded nanoemulsions (1) after frozen storage; (2) across freeze-thaw cycles; (3) after thawing;
vi. Figure S6: Modeling of blood-pool kinetics of nanoemulsions after bolus administration;
vii. Figure S7: Blood flow velocity changes after ultrasonic nicardipine uncaging in the aorta;
viii. Tables S1-S3: Tables summarizing plasma and whole blood half-lives of nanoemulsions and ultrasonically uncaged model drugs.
ix. Supplementary Videos S1 and S2: Videos demonstrating the distension of abdominal aorta upon ultrasonic nicardipine uncaging.

**Abbreviations**
FUS: focused ultrasound
PFP: perfluoropentane
PEG: polyethylene glycol
PLGA: poly(*D,L* lactic-co-glycolic acid) di-block copolymer
PCL: poly-ε-caprolactone
PLLA: poly-*L*-lactide
LAL: Limulus, Amebocyte Lysate
PDI: polydispersity
MRI: magnetic resonance imaging, i.v.: intravenous

## Acknowledgements

This work was funded with grant support from the National Institutes of Health (BRAIN Initiative RF1 MH114252), the Stanford Center for Cancer Nanotechnology Excellence, the Foundation of the American Society for Neuroradiology, and the Wallace H. Coulter Foundation. We would also like to thank William T. Newsome, PhD and the Stanford Neurosciences Institute for additional funding support. We would like to thank Yun-Sheng Chen, PhD for assistance in the design and preparation of the nanoemulsions, and Marko Jakovljevic, PhD and Jeremy Dahl, PhD for access to and training on the Siemens Acuson S2000 scanner. We would like to thank Kim Butts Pauly, PhD, Katherine Ferrara, PhD, Jennifer Dionne, PhD, and the whole Airan Lab for helpful discussions.

## Author Contributions

QZ and RDA designed the experiments. QZ performed the chemistry of nanoparticle production and physicochemical characterization. QZ performed the *in vitro/in vivo* ultrasonic drug uncaging experiments, with guidance from MA and assistance from JBW and AK. QZ and BCY performed the ultrasound imaging and ultrasonic nicardipine uncaging experiments and their analysis. QZ, BCY, and RDA prepared the figures and wrote the manuscript.

